# The transcriptional correlates of divergent electric organ discharges in *Paramormyrops* electric fish

**DOI:** 10.1101/625475

**Authors:** Mauricio Losilla, Jason R. Gallant

## Abstract

**Background:** Understanding the genomic basis of phenotypic diversity can be greatly facilitated by examining adaptive radiations with hypervariable traits. In this study, we focus on a rapidly diverged species group of mormyrid electric fish in the genus *Paramormyrops*, which are characterized by extensive phenotypic variation in electric organ discharges (EODs). The main components of EOD diversity are waveform duration, complexity and polarity. Using an RNA-sequencing based approach, we sought to identify gene expression correlates for each of these EOD waveform features by comparing 11 specimens of *Paramormyrops* that exhibit variation in these features.

**Results:** Patterns of gene expression among *Paramormyrops* are highly correlated, and 3,274 genes (16%) were differentially expressed. Using our most restrictive criteria, we detected 71-144 differentially expressed genes correlated with each EOD feature, with little overlap between them. The predicted functions of several of these genes are related to extracellular matrix, cation homeostasis, lipid metabolism, and cytoskeletal and sarcomeric proteins. These genes are of significant interest given the known morphological differences between electric organs that underlie differences in the EOD waveform features studied.

**Conclusions:** In this study, we identified plausible candidate genes that may contribute to phenotypic differences in EOD waveforms among a rapidly diverged group of mormyrid electric fish. These genes may be important targets of selection in the evolution of species-specific differences in mate-recognition signals.

## Introduction

Understanding the genomic basis of phenotypic diversity is a major goal of evolutionary biology [1]. Adaptive radiations and explosive diversification of species [2] are frequently characterized by interspecific phenotypic differences in divergence of few, hypervariable phenotypic traits [3–6]. Such systems offer exceptional advantages to study the genomic bases of phenotypic diversity: they can provide replication under a controlled phylogenetic framework [7], and couple ample phenotypic differentiation with relatively “clean” genomic signals between recently diverged species [8]. Study of the genomic mechanisms underlying hypervariable phenotypic traits has identified, in some cases relatively simple genetic architectures [9–13]. More often, the genetic architecture underlying such traits can be complex and polygenic [14–17]. It has long been recognized that changes in gene expression can affect phenotypic differences between species [18], and RNA-seq based approaches have greatly facilitated the study of this relationship [19]. A growing number of studies have examined differences in gene expression in phenotypic evolution (e.g., [19–27]). While these studies do not implicate mutational causes, analysis of differential gene expression (DGE) can be a useful approach in examining the genomic basis of divergent phenotypes.

African weakly electric fish (Teleostei: Mormyridae) are among the most rapidly speciating groups of ray-finned fishes [28,29]. This is partly due to the diversification of the genus *Paramormyrops* [30,31] in the watersheds of West-Central Africa, where more than 20 estimated species [32] have evolved within the last 0.5-2 million years [30]. Extensive evidence has demonstrated that electric organ discharges (EODs) exhibit little intraspecific variation, yet differ substantially among mormyrid species [33–35]. This pattern is particularly evident in *Paramormyrops* [30,36], in which EOD waveforms evolve much faster than morphology, size, and trophic ecology [37].

Mormyrid EODs are a behavior with a dual role in electrolocation [38,39] and intraspecific communication [40,41]. EOD waveforms vary between species principally in terms of their complexity, polarity, and duration [30,42], and all three dimensions of variation are evident among *Paramormyrops* (Fig. 1). Furthermore, recent discoveries of intraspecific polymorphism in EOD waveform in *P. kingsleyae* [43] and polarity among *P.* sp. ‘magnostipes’ [35] present a unique opportunity to study the genomic basis of phenotypic traits within a rapidly diverging species group.

**Figure 1.**
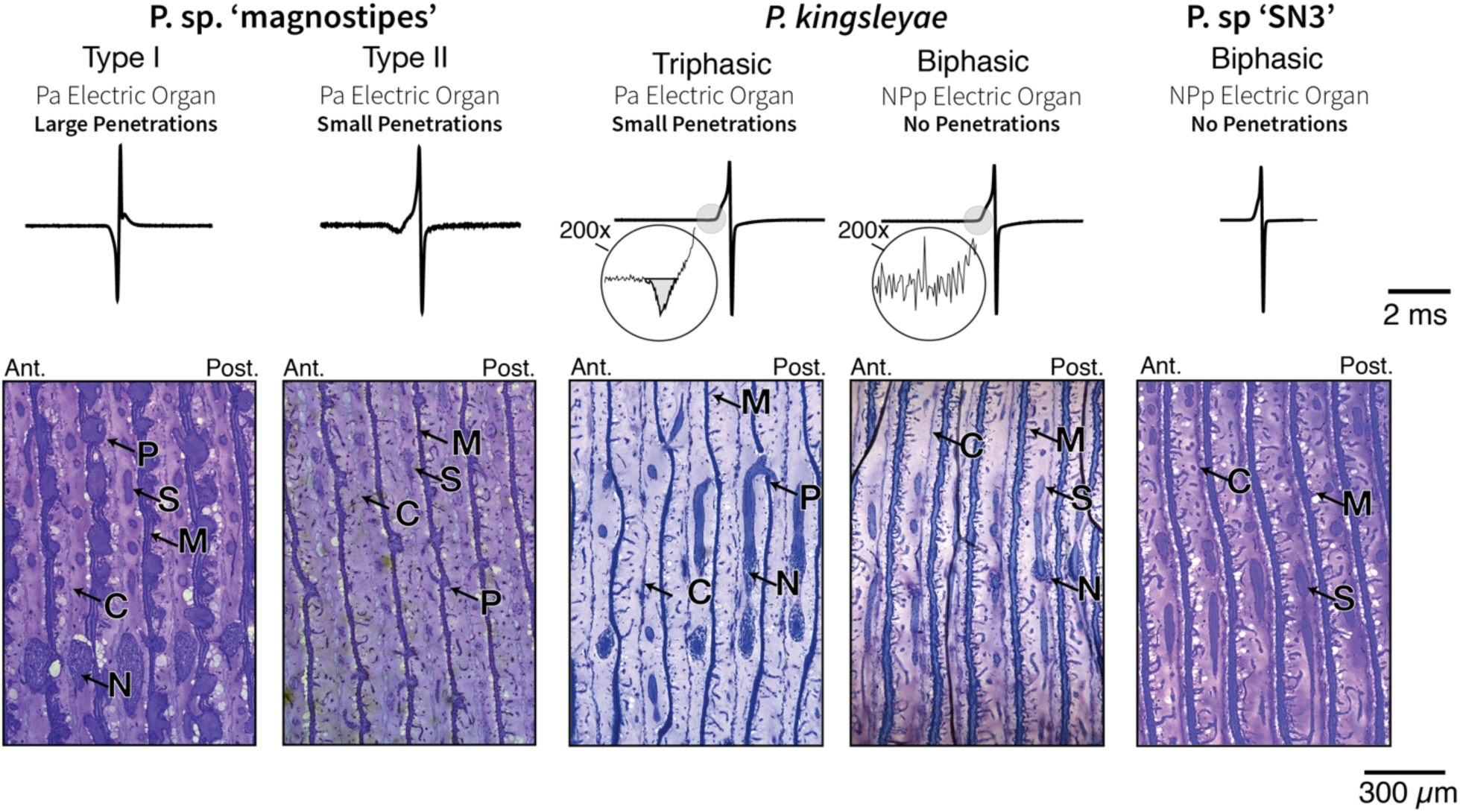
Electric organ discharge (EOD) diversity and electric organ anatomy in *Paramormyrops*. EOD traces from specimens in this study and representative parasagittal sections of the five *Paramormyrops* operational taxonomic units (OTUs) considered in this study. 200x magnification on *P. kingsleyae* EODs reveals a P0 phase on triphasic EODs only. Individuals with triphasic EODs all have penetrations, whereas individuals with biphasic EODs do not. OTUs with ‘inverted’ polarity triphasic EODs have large penetrations compared to OTUs with normal polarity triphasic EODs. Ant. = anterior, C = connective tissue septa, N = nerve, M = microstalklets (profusely branched stalks), P = penetrations, Post. = posterior, S = stalks.

EODs have a well-understood morphological (Fig. 1) and neurophysiological basis [44,45]. EODs are generated by specialized cells (electrocytes) that constitute the electric organ (EO), located in the caudal peduncle [46]. Mormyrid EOs are comprised of 80-360 electrocytes [34], and an individual EOD is produced when the electrocytes discharge synchronously. EODs are multiphasic because they result from action potentials produced by two excitable membranes: the two large phases of the EOD, called P1 and P2, are produced by spikes generated by the posterior and anterior electrocyte faces, respectively [47]. There is a relationship between EODs of longer duration and increased surface membrane area [48], likely mediated at least in part by an increase in membrane capacitance [49,50]. The duration of EODs is highly variable within mormyrids-- some EODs are extremely long (>15 ms) and others are very brief (0.2 ms) [32].

Within the Mormyridae, triphasic EODs evolved early from biphasic EODs; however, there have been multiple parallel reversions to biphasic EODs across mormyrids and within *Paramormyrops* [36,43]. Triphasic (P0-present) EODs are produced by electrocytes that are innervated on the anterior face and have penetrating stalks (*Pa*, P-type), whereas biphasic (P0-absent) EODs are produced by electrocytes innervated on the posterior face and lack penetrating stalks (*NPp*, N-type) (for more details see [42,43,47,48,51,52]). We refer to triphasic EODs as more ‘complex’ than biphasic EODs. In some cases, triphasic EODs display an unusually large P0 phase, which gives the appearance of an ‘inverted’ polarity. This is exemplified by the type I EODs of *P.* sp. ‘magnostipes’ (Fig. 1) [35]. The number [47] and diameter [34,43] of stalk penetrations are positively correlated with the magnitude of P0. We refer to individuals with large penetrations as ‘inverted’ polarity and individuals with small penetrations as ‘normal’ polarity.

Recent studies in mormyrids [53–57] have adopted a candidate gene approach to examine the molecular basis of variation in EOD duration on macroevolutionary scales, implicating voltage gated sodium channels (e.g. *scn4aa*) and potassium channels (e.g. *kcna7a*) as key targets of selection during EOD evolution. Beyond this recent attention to ion channels, several studies have described the importance of structural differences between EOs as an important component of EOD variation [43,48,50]. In this study, we took a transcriptome-wide approach to characterizing the molecular basis of electric signal diversity in *Paramormyrops* species divergent for EOD complexity, duration and polarity. We used RNA-sequencing to comprehensively examine DGE in the adult EOs of five *Paramormyrops* operational taxonomic units (OTUs), leveraging a recently sequenced and annotated genome assembly from the species *P. kingsleyae* (N-type) [58], and identify gene expression correlates of each of the three main EOD waveform features of electric signal diversity in *Paramormyrops*. Our results emphasize genes that influence the shape and structure of the electrocyte cytoskeleton, membrane and extracellular matrix (ECM) to exhibit predictable differences between *Paramormyrops* species with divergent EOD phenotypes.

## Methods

### Sample collection

We captured 11 *Paramormyrops* individuals from Gabon, West Central Africa in 2009: five *P. kingsleyae* (n=3 N-type and n=2 P-type), four *P.* sp. ‘magnostipes’ (n=2 Type I and n=2 Type II), and two *P.* sp. ‘SN3’. Within 1-12 hours of capture, individual specimens were euthanized by overdose with MS-222. The caudal peduncle was excised and skinned, and immediately immersed in RNA-later for 24h at 4°C, before being transferred to −20°C for long-term storage. As two of these species (*P.* sp. ‘magnostipes’, *P.* sp. ‘SN3’) are presently undescribed, we note that these specimens were identified by their EOD waveform, head morphology and collecting locality [30,31,35,59]. All specimens, including vouchers materials, are deposited in the Cornell University Museum of Vertebrates. Collection information and the phenotypes per EOD feature of each sample are detailed in Table 1.

**Table 1.** Phenotypic and collection information of the samples studied.

| Tag No. | OTU | Phenotypes per EOD feature |  |  | CUMV<br>Accession<br>Number | Site Name | Lat,<br>Long | SL<br>(mm) | Sex |
| --- | --- | --- | --- | --- | --- | --- | --- | --- | --- |
|  |  | Duration | Complexity | Polarity (diameter of<br>penetrations) |  |  |  |  |  |
| PKINGN_6898 | *P. kingsleyae* (N-type) | long EOD | biphasic | NA (no penetrations) | 95184 | Bikagala<br>Creek | -2.20,<br>11.56 | 131 | M |
| PKINGN_6900 | *P. kingsleyae* (N-type) | long EOD | biphasic | NA (no penetrations) | 95184 | Bikagala<br>Creek | -2.20,<br>11.56 | 145 | M |
| PKINGN_6901 | *P. kingsleyae* (N-type) | long EOD | biphasic | NA (no penetrations) | 95184 | Bikagala<br>Creek | -2.20,<br>11.56 | 149 | M |
| PKINGP_6716 | *P. kingsleyae* (P-type) | long EOD | triphasic | small penetrations | 95183 | Mouvanga<br>Creek | -2.33,<br>11.69 | 92.5 | J/F |
| PKINGP_6718 | *P. kingsleyae* (P-type) | long EOD | triphasic | small penetrations | 95183 | Mouvanga<br>Creek | -2.33,<br>11.69 | 91 | J/F |
| PMAG1_6780 | *P. sp.* 'magnostipes type I' | long EOD | triphasic | large penetrations | 95155 | Mouvanga<br>Creek | -2.33,<br>11.69 | 107 | M |
| PMAG1_6787 | *P. sp.* 'magnostipes type I' | long EOD | triphasic | large penetrations | 95155 | Mouvanga<br>Creek | -2.33,<br>11.69 | 97.5 | F |
| PMAG2_6768 | *P. sp.* 'magnostipes type II' | long EOD | triphasic | small penetrations | 95155 | Mouvanga<br>Creek | -2.33,<br>11.69 | 93 | M |
| PMAG2_6769 | *P. sp.* 'magnostipes type II' | long EOD | triphasic | small penetrations | 95155 | Mouvanga<br>Creek | -2.33,<br>11.69 | 124 | M |
| PSN3_6739 | *P. sp.* 'SN3' | short EOD | biphasic | NA (no penetrations) | Uncatalogued | Mouvanga<br>Creek | -2.33,<br>11.69 | 73 | J/F |
| PSN3_6742 | *P. sp.* 'SN3' | short EOD | biphasic | NA (no penetrations) | 95173 | Mouvanga<br>Creek | -2.33,<br>11.69 | 70 | J/F |

CUMV = Cornell University Museum of Vertebrates, EOD = electric organ discharge, F= female, J = juvenile, Lat = Latitude, Long = Longitude, M = male, NA = not applicable, OTU = operational taxonomic unit, SL = standard length

### RNA extraction, cDNA library preparation and Illumina Sequencing

Total RNA was extracted from EOs using RNA-easy Kit (Qiagen, Inc) after homogenization with a bead-beater (Biospec, Inc.) in homogenization buffer. mRNA was isolated from total RNA using a NEBNext mRNA Isolation Kit (New England Biolabs, Inc.). Libraries for RNA-seq were prepared using the NEBNext mRNA Sample Prep Master Mix Set, following manufacturer’s instructions. Final libraries after size selection ranged from 250-367 bp. Libraries were pooled and sequenced by the Cornell University Biotechnology Resource Center Genomics Core on an Illumina HiSeq 2000 in a 2×100bp paired end format. Raw sequence reads were deposited in the NCBI SRA (Supplemental File 1).

### Read processing and data exploration

FastQC v0.11.3 (Babraham Bioinformatics) was used to manually inspect raw and processed reads. We used Trimmomatic v.0.32 [60] to remove library adaptors, low quality reads, and filter small reads; following the suggested settings of MacManes [61]: 2:30:10 SLIDINGWINDOW:4:5 LEADING:5 TRAILING:5 MINLEN:25. After trimming, reads from each specimen were aligned to the predicted transcripts of the NCBI-annotated (Release 100) *P. kingsleyae* (N-type) genome [58] using bowtie 2 v2.3.4.1 [62]. Expression quantification was estimated at the gene level using RSEM v1.3.0 [63], followed by exploration of the data with a gene expression correlation matrix based on Euclidean distances and Pearson’s correlation coefficient (for genes with read counts >10, Trinity’s default parameters). All these steps were executed using scripts included with Trinity v2.6.6 [64,65].

### Data Analysis

We began by examining DGE between all possible pairwise comparisons of OTUs (n = 10, Table 2) using edgeR v3.20.9 [66] through a script provided with Trinity. We restricted our consideration of genes to those where CPM-transformed counts were > 1 in at least two samples for each comparison (edgeR default parameters). We modified this to use the function estimateDisp() instead of the functions estimateCommonDisp() and estimateTagwiseDisp(). For each comparison, we conservatively considered genes to be differentially expressed with a minimum fold change of 4 and p-value of 0.001 after FDR correction. We compiled a non-redundant list of genes that were differentially expressed in at least one comparison based on these criteria (Fig. 2, Set A).

**Table 2.** All ten possible pairwise DGE comparisons with the total number of DEG and enriched GO terms for each. Also indicated is whether each comparison is informative for contrasting each EOD feature. The phenotypes for waveform polarity can only be contrasted in comparisons where both OTUs have penetrations. Informative comparisons for each EOD feature (Set A’) are marked with an * in the column of the EOD feature they contrasted.

| Comparison |  | Contrast |  |  | DEG genes | enriched GO terms |  |  |
| --- | --- | --- | --- | --- | --- | --- | --- | --- |
| OTU #1 | OTU #2 | Duration | Complexity | Polarity |  | BP | CC | MF |
| *P. kingsleyae* (N-type) | *P. kingsleyae* (P-type) | no | yes* | NA (no) | 530 | 46 | 12 | 17 |
| *P. kingsleyae* (N-type) | *P. sp.* 'magnostipes type I' | no | yes | NA (no) | 1542 | 76 | 15 | 37 |
| *P. kingsleyae* (N-type) | *P. sp.* 'magnostipes type II' | no | yes | NA (no) | 1174 | 71 | 16 | 25 |
| *P. kingsleyae* (N-type) | *P. sp.* 'SN3' | yes* | no | NA (no) | 507 | 52 | 4 | 13 |
| *P. kingsleyae* (P-type) | *P. sp.* 'magnostipes type I' | no | no | yes* | 1322 | 40 | 12 | 25 |
| *P. kingsleyae* (P-type) | *P. sp.* 'magnostipes type II' | no | no | no | 719 | 47 | 10 | 24 |
| *P. kingsleyae* (P-type) | *P. sp.* 'SN3' | yes | yes | NA (no) | 385 | 33 | 3 | 14 |
| *P. sp.* 'magnostipes type I' | *P. sp.* 'magnostipes type II' | no | no | yes | 9 | 5 | 1 | 1 |
| *P. sp.* 'magnostipes type I' | *P. sp.* 'SN3' | yes | yes | NA (no) | 1053 | 43 | 6 | 27 |
| *P. sp.* 'magnostipes type II' | *P. sp.* 'SN3' | yes | yes | NA (no) | 489 | 40 | 9 | 16 |

BP = biological process, CC = cellular component, DEG = differentially expressed genes, DGE = differential gene expression, GO = gene ontology, NA = not applicable, MF = molecular function, OTU = operational taxonomic unit

**Figure 2.**
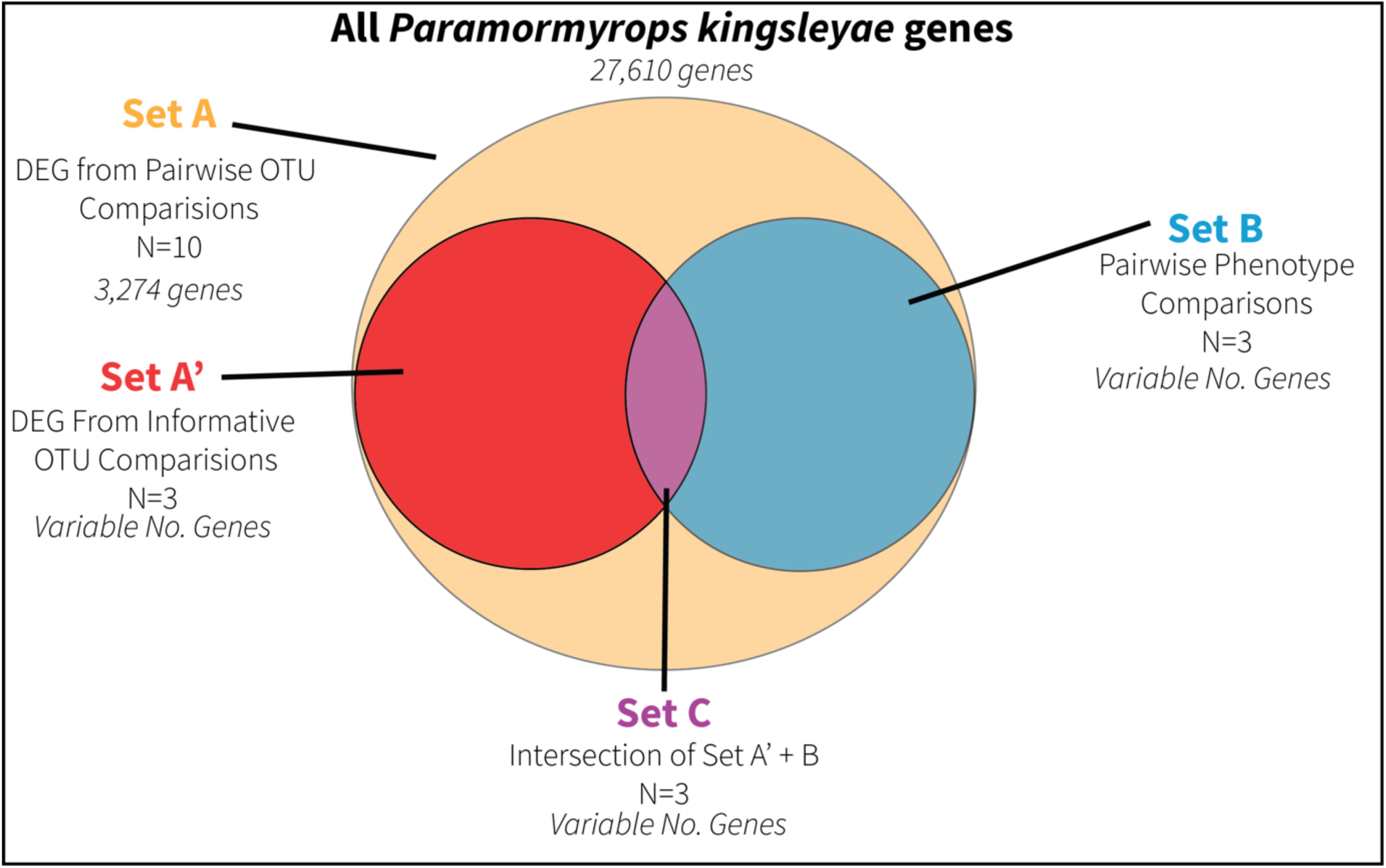
Diagram of how we constructed the lists of upregulated genes of Set C. We used the same approach to find the respective enriched GO terms. DEG = differentially expressed genes, N = number of comparisons made for each set, OTU = operational taxonomic unit.

For each of the differentially expressed genes (DEGs) in Set A, we TMM normalized, log2(TMM +1) transformed, and mean-centered their expression values. We used the transformed values to compare gene expression between groups of OTUs with alternative EOD waveform phenotypes (i.e. *long duration EOD vs. short duration EOD*, *biphasic vs. triphasic* and *small penetrations vs. large penetrations,* see Table 1. Note that waveform polarity phenotypes only apply to triphasic individuals). For each of the three phenotype pairs, we extracted the genes that were on average more than four times more highly expressed in one phenotype than the other. This resulted in six lists of upregulated genes, one for each EOD feature across all OTUs and samples (Fig. 2, Set B).

In order to assess enrichment of particular gene pathways, biological functions, and cellular locations using a controlled vocabulary, we performed Gene Ontology (GO) [67,68] enrichment tests on every list of upregulated genes from (1) the ten pairwise comparisons (n=20, two per comparison) and (2) Set B (n=6), for each of the three ontology domains: Biological Process, Cellular Component, and Molecular Function. First, we identified homologous proteins predicted from the *P. kingsleyae* (N-type) reference genome and those predicted from *Danio rerio* (GRCz11) by blastp (BLAST+ v2.6, [69]). For each protein, the top hit (e-value ≤ 1e-10) was used for annotation. Next, we used mygene v1.14.0 [70,71] to match the *D. rerio* proteins to *D. rerio* genes and extract their GO annotations (zebrafish Zv9). This resulted in GO annotations for each of the three ontology domains for *P. kingsleyae* (N-type) genes. Finally, we carried out the GO enrichment tests using topGO v2.30.1 [72] and the following parameters: nodeSize = 10, statistic = fisher, algorithm = weight01, p-value ≤ 0.02. The ‘universe’ for each enrichment test on gene lists from the pairwise comparisons was all the genes deemed expressed in the respective comparison, whereas the non-redundant list of genes in these ten ‘universes’ was the ‘universe’ for all enrichment tests on the gene lists from Set B.

Interpretation of lists of genes from Set A and Set B each suffered limitations for the overall goals of this analysis, which is to identify the DEGs most strongly associated with each waveform feature (duration, complexity, and polarity). The ten comparisons made to construct Set A were not equally informative for two primary reasons: (1) the OTUs in this analysis vary in terms of their phylogenetic relatedness (see [30,31]) and (2) several OTU comparisons varied in more than one waveform characteristic (Table 2). As such, we elected to focus on the most informative comparison for each EOD feature: the comparison that contrasted only the given feature and that minimized phylogenetic distance between OTUs. Of the ten pairwise comparisons, we classified three as the most informative comparisons, one per EOD feature (Table 2). The six lists of upregulated genes from these three comparisons constitute Set A’.

Comparisons in Set A’; however, lack biological replication. In contrast, interpretations of Set B were potentially limited in that many of the OTUs in this analysis differed in more than one EOD feature. To circumvent the limitations of Sets A’ and B within the limits of our study design, we constructed a third set (Set C). Set C is defined as the intersection of the upregulated genes and their enriched GO terms from Sets A’ and B, for each phenotype. Since there were six phenotypes in our study, Set C encompasses six lists of upregulated genes and their respective enriched GO terms (Fig. 2). Therefore, Set C represents the DEGs that are (1) differentially expressed between closely-related OTUs that vary in a single waveform characteristic, and (2) are consistently differentially expressed among all OTUs that share that waveform feature. We focus our attention on Set C: We retrieved GO term definitions from QuickGO [73] and descriptions of gene function of the functional annotations from UniProt [74]; and to facilitate the discussion, we classified the more interesting genes in Set C into “general” functional classes, or themes. All source code necessary to perform the methods described here is provided in a GitHub repository: http://github.com/msuefishlab/paramormyrops_rnaseq.

## Results

### Overall Results

Overall alignment rates to the *Paramormyrops kingsleyae* reference transcriptome ranged from 28-74% (>375 million sequenced reads in total, 50% aligned), with no clear differences among OTUs (Supplemental File 1). On inspection, we concluded that these rates are a consequence of the presence of overrepresented sequences from rRNA, mtDNA and bacterial contamination in the RNA-seq reads.

Fig. 3 shows a heatmap of pairwise correlations of gene expression for 24,960 genes across all 11 samples. Our ten DGE comparisons detected a range of 16,420-19,273 expressed genes. Intersection of these lists resulted in a non-redundant list of 20,197 genes expressed in EO across all DGE comparisons. We found that 3,274 (16%) were differentially expressed in at least one comparison, and expression patterns across all OTUs were highly correlated (Pearson’s r > 0.89, Fig. 3). Despite this, correlation values were higher among recognized OTUs, except for the *P.* sp. ‘magnostipes type II’ 6768 sample (Fig. 3). Thus, we excluded comparisons with the *P.* sp. ‘magnostipes type II’ OTU from the informative comparisons for Set A’.

**Figure 3.**
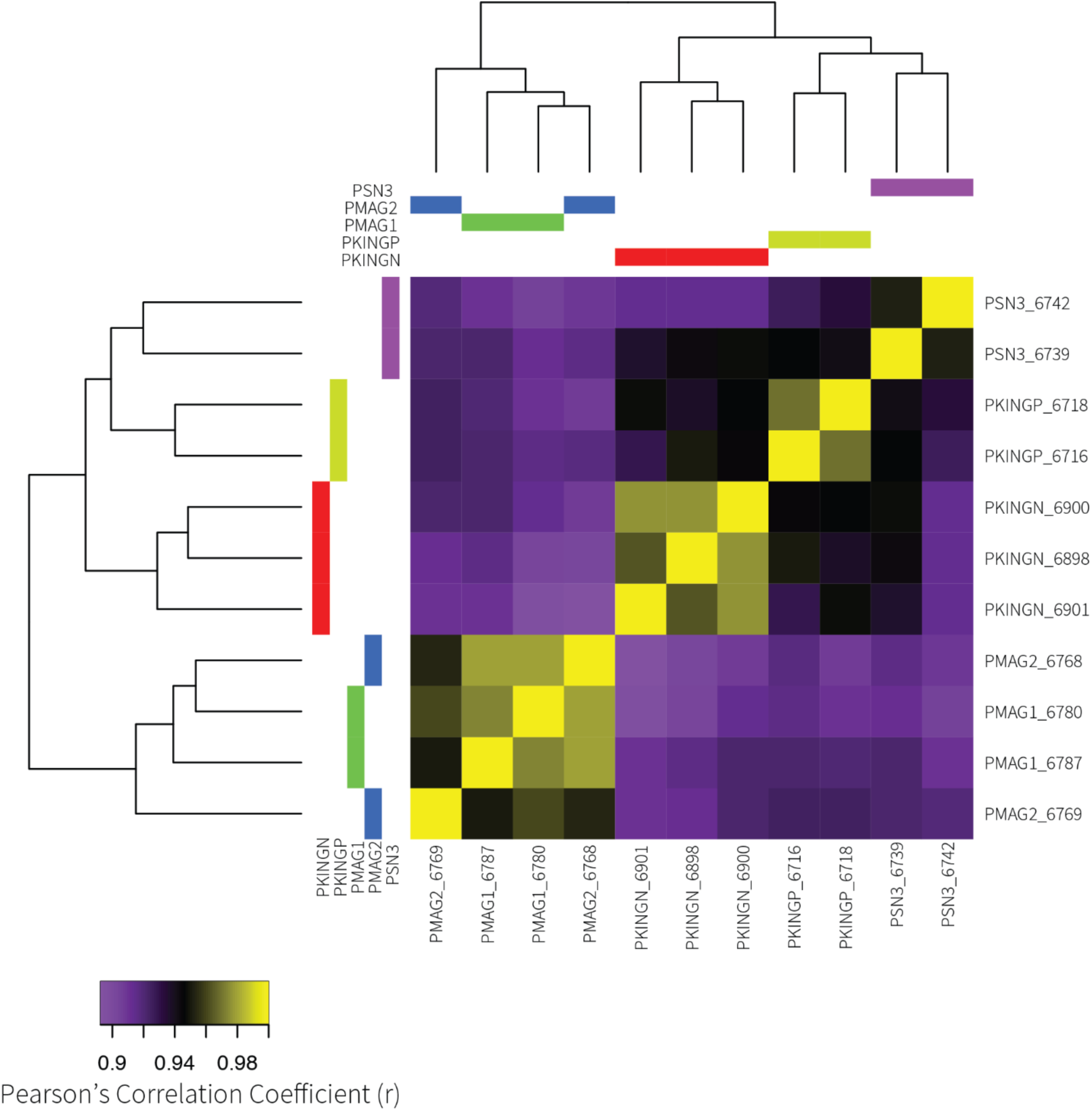
Heatmap of sample by sample correlations in gene expression, and the inferred phylogenetic relationships from these expression correlation values.

### Set A: Differential Expression Analysis

We found between 489-1542 DEGs (50-128 enriched GO terms) in every comparison except *P.* sp. ‘magnostipes type I’ vs *P.* sp. ‘magnostipes type II’, which had only nine DEGs with seven enriched GO terms (Table 2). Supplemental File 2 provides a tabular list of DEGs for each comparison, and Supplemental File 3 provides a tabular list of enriched GO terms for each comparison.

We chose the phylogenetically most informative comparisons (see methods) to construct Set A’, which are indicated in Table 2. We found: 507 DEG and 69 enriched GO terms comparing *P. kingsleyae* (N-type) vs *P.* sp. ‘SN3’ (EOD duration); 1322 DEG and 77 enriched GO terms comparing *P. kingsleyae* (P-type) vs *P.* sp. ‘magnostipes type I’ (waveform polarity); and 530 DEG and 75 enriched GO terms comparing *P. kingsleyae* (N-type) vs *P. kingsleyae* (P-type) (waveform complexity).

### Set B: Expression Based Clustering

For each EOD feature (n=3), we grouped OTUs by phenotype (Table 1), and calculated normalized expression values for Set A genes (n = 3,274). For each EOD feature, we selected genes exhibiting a greater than four-fold difference in averaged, normalized expression between phenotypes to construct Set B. The expression profiles of the genes in the clusters for each EOD feature, along with the enriched GO terms for Biological Process and Cellular Component, are shown in Figs. 4–6. Supplemental File 4 lists the identities of these DEG and Supplemental File 5 lists their enriched GO terms for all three GO ontologies.

**Figure 4.**
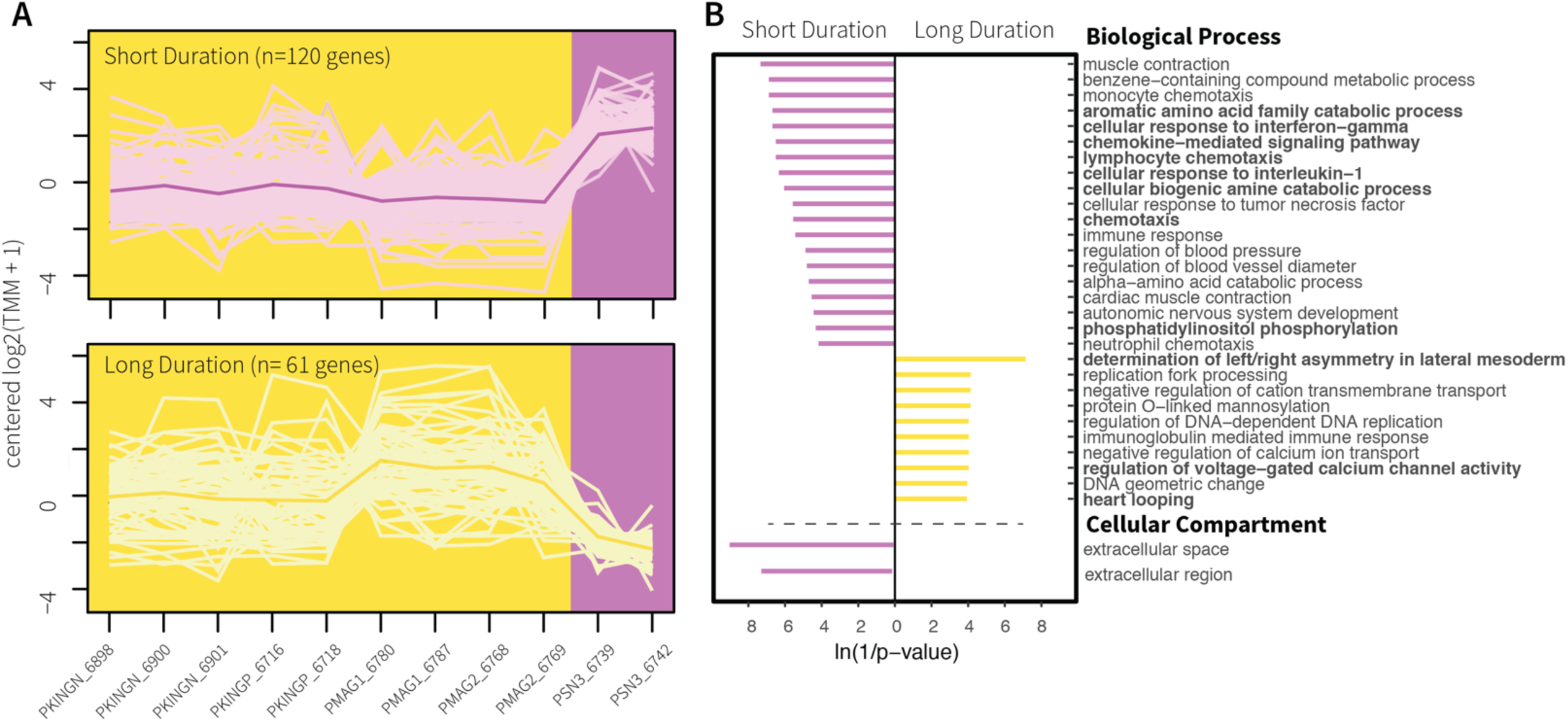
A) Gene clusters whose expression patterns correlate with the electric organ discharge (EOD) duration phenotypes short EODs (purple background) and long EODs (yellow background). Samples are sorted alphabetically on the X axis. The lines connect transformed gene expression values across all samples; light-color lines represent one gene, the dark-color line is the average expression pattern of all genes in the cluster. B) Gene Ontology (GO) terms for Biological Process and Cellular Component found enriched in the gene clusters from (A). The X axis shows transformed p-values, the longer a bar the smaller its p-value. The direction and color of a bar indicate the phenotype in which the GO term is enriched [same color code as (A)]. GO terms highlighted in bold also belong to Set C.

**Figure 5.**
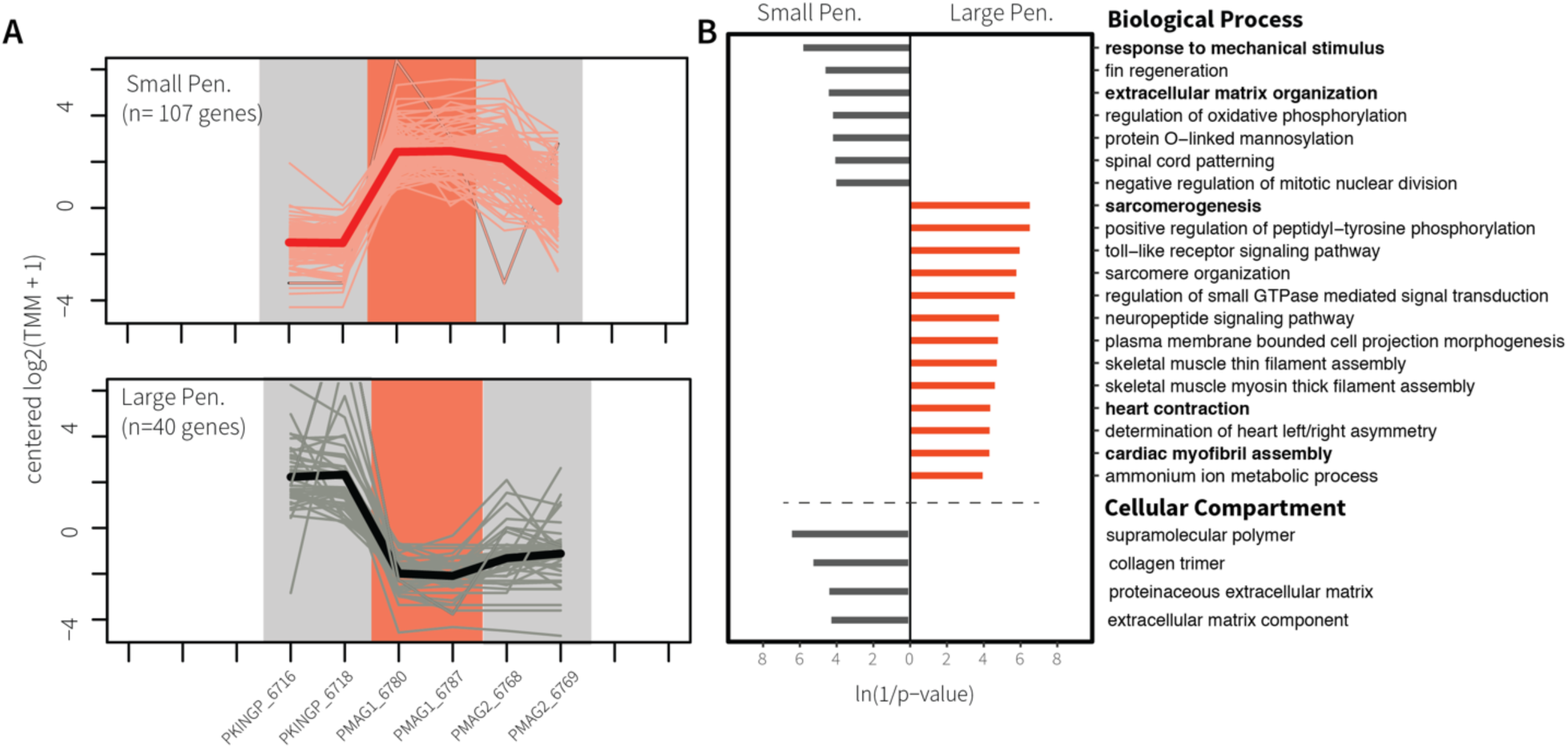
A) Gene clusters whose expression patterns correlate with the electric organ discharge (EOD) waveform polarity phenotypes small penetrations (grey background) and large penetrations (salmon background). Samples are sorted alphabetically on the X axis, but only samples with penetrations are considered for this EOD feature. The lines connect transformed gene expression values across all samples; light-color lines represent one gene, the dark-color line is the average expression pattern of all genes in the cluster. B) Gene Ontology (GO) terms for Biological Process and Cellular Component found enriched in the gene clusters from (A). The X axis shows transformed p-values, the longer a bar the smaller its p-value. The direction and color of a bar indicate the phenotype in which the GO term is enriched [same color code as (A)]. GO terms highlighted in bold also belong to Set C. Pen = penetrations.

**Figure 6.**
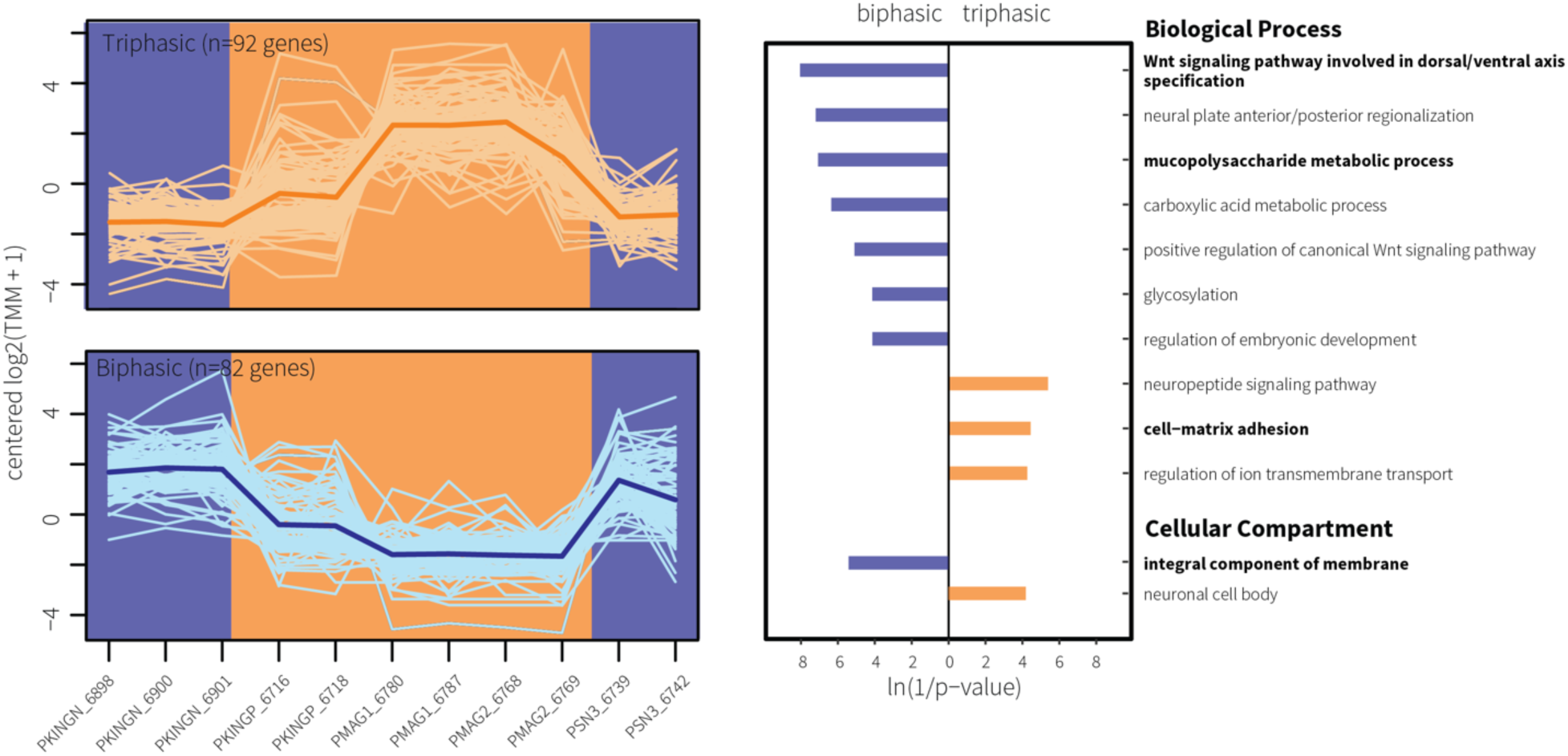
A) Gene clusters whose expression patterns correlate with the electric organ discharge (EOD) waveform complexity phenotypes biphasic (blue background) and triphasic (orange background). Samples are sorted alphabetically on the X axis. The lines connect transformed gene expression values across all samples; light-color lines represent one gene, the dark-color line is the average expression pattern of all genes in the cluster. B) Gene Ontology (GO) terms for Biological Process and Cellular Component found enriched in the gene clusters from (A). The X axis shows transformed p-values, the longer a bar the smaller its p-value. The direction and color of a bar indicate the phenotype in which the GO term is enriched [same color code as (A)]. GO terms highlighted in bold also belong to Set C.

Contrast of waveform duration identified 181 DEG and 40 enriched GO terms. 121 of the DEG were upregulated in the short EOD phenotype (Fig. 4A, purple lines). These genes were enriched with GO terms that include ‘extracellular space,’ ‘extracellular region,’ ‘muscle contraction,’ and ‘phosphatidylinositol phosphorylation’ (Fig. 4B, purple bars), whereas 61 genes were upregulated in samples with long EODs (Fig. 4A, yellow lines). These genes were enriched with GO terms like ‘negative regulation of cation transmembrane transport,’ ‘regulation of voltage-gated calcium channel activity,’ and ‘neuropeptide hormone activity’ (Fig. 4B, yellow bars).

Contrast of waveform polarity identified 147 DEG and 35 enriched GO terms. We found 40 upregulated genes (Fig. 5A, grey lines) in individuals with small penetrations. These genes were enriched with GO terms such as ‘response to mechanical stimulus,’ ‘extracellular matrix organization,’ and ‘collagen trimer’ (Fig. 5B, grey bars). In large penetrations phenotype, we found 107 upregulated genes (Fig. 5A, salmon lines). These genes were enriched with GO terms that included ‘sarcomere organization,’ ‘plasma membrane bounded cell projection morphogenesis,’ ‘calcium ion binding,’ and ‘structural constituent of cytoskeleton’ (Fig. 5B, salmon bars).

Finally, contrast of waveform complexity identified 174 DEG and 16 enriched GO terms. We detected 82 upregulated genes in individuals with biphasic EODs (Fig. 6A, blue lines). These genes were enriched with GO terms like ‘positive regulation of canonical Wnt signaling pathway,’ ‘mucopolysaccharide metabolic process,’ and ‘integral component of membrane’ (Fig. 6B, blue bars). We detected 92 upregulated genes in individuals with triphasic EODs (Fig. 6A, orange lines). These genes were enriched with GO terms that included ‘cell-matrix adhesion,’ ‘regulation of ion transmembrane transport,’ and ‘neuronal cell body’ (Fig. 6B, orange bars).

### Set C: Intersection of Phylogenetically Informative Comparisons and Expression Based Clustering

We were motivated to obtain the DEGs and enriched GO terms that were most likely to be associated with divergent EOD phenotypes. To obtain this list, we constructed Set C, which is the intersection of Set A’ and Set B described above.

Contrast of waveform duration identified 105 DEG and 11 enriched GO terms. 79 of the DEG were upregulated in the short EOD phenotype, and 26 genes were upregulated in samples with long EODs. Contrast of waveform polarity identified 144 DEG and 9 enriched GO terms. We found 38 upregulated genes in individuals with small penetrations and 106 upregulated genes in individuals with large penetrations. Finally, contrast of waveform complexity identified 71 DEG and 4 enriched GO terms. We detected 46 upregulated genes in individuals with biphasic EODs and 25 upregulated genes in individuals with triphasic EODs. These results are further detailed in Table 3. The enriched GO terms in Set C for Biological Process and Cellular Component are emphasized in boldface for waveform duration (Fig. 4B), polarity (Fig. 5B), and complexity (Fig. 6B). The individual genes in Set C are listed in Supplemental File 6 and their associated GO terms are listed in Supplemental File 7.

**Table 3.** Total number of upregulated genes and enriched GO terms in Set C for each EOD feature, phenotype and ontology.

| EOD feature | Phenotype | Upregulated genes | Protein-coding (%) | enriched GO terms |  |  |
| --- | --- | --- | --- | --- | --- | --- |
|  |  |  |  | BP | CC | MF |
| Duration | short EODs | 79 | 65 (82) | 8 | 0 | 0 |
| Duration | long EODs | 26 | 21 (81) | 3 | 0 | 0 |
| Polarity | small penetrations | 38 | 28 (74) | 2 | 0 | 1 |
| Polarity | large penetrations | 106 | 87 (82) | 3 | 0 | 3 |
| Complexity | biphasic | 46 | 34 (74) | 2 | 1 | 0 |
| Complexity | triphasic | 25 | 23 (92) | 1 | 0 | 0 |

BP = biological process, CC = cellular component, EOD = electric organ discharge, GO = gene ontology, MF = molecular function.

## Discussion

It has long been recognized that changes in gene expression can affect phenotypic differences between species [18], and RNA-seq has facilitated the study of this relationship [19]. The goal of this study was to determine DEGs associated with divergent EOD features within *Paramormyrops*. Expression patterns across all OTUs were highly correlated (Pearson’s r > 0.89, Fig. 3) and we detected differential expression of only 3,274 (16%) genes between any two OTUs. Thus, a major finding of this study is that EO gene expression is overall quite similar across *Paramormyrops* species with divergent EODs, and relatively few genes are associated with phenotypic differences in EOD waveform between OTUs. Given generally high levels of genetic distances observed between geographically proximate populations of these *Paramormyrops* species [35,75] this was a somewhat unexpected finding.

Despite the relatively small number of DEGs compared to the total number of genes expressed in the EO, we constructed our analysis to extract genes that were highly associated with particular phenotypes. Set A’ represents a formal statistical test that contrasted OTUs. Each comparison contrasted samples from OTUs that were divergent in only one EOD feature, while minimizing phylogenetic distance. The tradeoff of this approach is small sample sizes without biological replication and potentially confounding variables, such as collection sites. Set B followed the opposite approach by minimizing the problem of biological replication at the expense of confounding phylogenetic relatedness and phenotypic heterogeneity. To mitigate this, we constructed Set C, which represents genes and GO terms that are differentially expressed/enriched between closely related OTUs divergent in only one phenotypic character and that are also consistently differentially expressed/enriched among representatives with similar EOD phenotypes. As such, we focus our discussion on the results of Set C. We classified the genes in Set C into “general” functional classes, or themes; and focus our attention on the ones that relate to the known morphological underpinnings of waveform duration (Table 4), polarity (Table 5), and complexity (Table 6). These functional classes were genes related to the ECM, cation homeostasis, lipid metabolism, and cytoskeletal and sarcomeric genes.

**Table 4.** Selected DEG in Set C for waveform duration by “general” functional class and EOD phenotype, and highlights of their predicted function.

| Pking_Entrez_geneID | Pking_gene_name | Pking_gene_symbol | “general” functional class | upregulated in phenotype | Highlights of Predicted Function (edited from UniProt) |
| --- | --- | --- | --- | --- | --- |
| 111832799 | *stathmin domain containing 1* | *stmnd1* | Cytoskeletal & sarcomeric | long EODs | GO MF: tubulin binding; GO BP: microtubule depolymerization, regulation of cytoskeleton organization |
| 111833088 | *myosin-7-like* | LOC111833088 | Cytoskeletal & sarcomeric | long EODs | Myosins are actin-based motor molecules with ATPase activity essential for muscle contraction |
| 111856289 | *pleckstrin homology-like domain family B member 1* | LOC111856289 | Cytoskeletal & sarcomeric | long EODs | GO BP: regulation of microtubule cytoskeleton organization |
| 111842483 | *parvalbumin-2* | LOC111842483 | Cytoskeletal & sarcomeric | short EODs | In muscle, parvalbumin is thought to be involved in relaxation after contraction. It binds two calcium ions |
| 111846153 | *troponin I, slow skeletal muscle-like* | LOC111846153 | Cytoskeletal & sarcomeric | short EODs | Inhibitory subunit of troponin, the thin filament regulatory complex which confers calcium-sensitivity to striated muscle actomyosin ATPase activity |
| 111856036 | *parvalbumin-2-like* | LOC111856036 | Cytoskeletal & sarcomeric | short EODs | In muscle, parvalbumin is thought to be involved in relaxation after contraction. It binds two calcium ions |
| 111860236 | *tropomyosin alpha-1 chain-like* | LOC111860236 | Cytoskeletal & sarcomeric | short EODs | Binds to actin filaments in muscle and non-muscle cells. Plays a central role, in association with the troponin complex, in the calcium dependent regulation of vertebrate striated muscle contraction. In non-muscle cells is implicated in stabilizing cytoskeleton actin filaments. |
| 111833641 | *hyaluronidase-5-like* | LOC111833641 | Extracellular matrix | short EODs | Catalyzes the hydrolysis of hyaluronan into smaller oligosaccharide fragments |
| 111837392 | *fibroblast growth factor binding protein 1* | *fgfbp1* | Extracellular matrix | short EODs | Acts as a carrier protein that releases fibroblast-binding factors (FGFs) from the extracellular matrix (EM) storage and thus enhance the mitogenic activity of FGFs |
| 111860877 | *collagenase 3-like* | LOC111860877 | Extracellular matrix | short EODs | Plays a role in the degradation of extracellular matrix proteins |
| 111834720 | *protein EFR3 homolog B-like* | LOC111834720 | Lipid metabolism | short EODs | Component of a complex required to localize phosphatidylinositol 4-kinase (PI4K) to the plasma membrane. The complex acts as a regulator of phosphatidylinositol 4-phosphate (PtdIns4P) synthesis |
| 111840357 | *PTB domain-containing engulfment adapter protein 1-like* | LOC111840357 | Lipid metabolism | short EODs | Modulates cellular glycosphingolipid and cholesterol transport |
| 111846286 | *retinoic acid receptor responder protein 3-like* | LOC111846286 | Lipid metabolism | short EODs | Catalyzes the calcium-independent hydrolysis of acyl groups in various phosphatidylcholines (PC) and phosphatidylethanolamine (PE) |
| 111847640 | *phosphatidylinositol 3-kinase regulatory subunit gamma-like* | LOC111847640 | Lipid metabolism | short EODs | Binds to activated (phosphorylated) protein-tyrosine kinases through its SH2 domain and regulates their kinase activity |
| 111852518 | *proto-oncogene c-Fos-like* | LOC111852518 | Lipid metabolism | short EODs | In growing cells, activates phospholipid synthesis, possibly by activating CDS1 and PI4K2A |

BP = biological process, DEG = differentially expressed genes, EOD = electric organ discharge, GO = gene ontology, MF = molecular function.

**Table 5.** Selected DEG in Set C for waveform polarity, by “general” functional class and EOD phenotype, and highlights of their expected function.

| Pking_Entrez_geneID | Pking_gene_name | Pking_gene_symbol | “general” functional class | upregulated in phenotype | Highlights of Predicted Function (edited from UniProt) |
| --- | --- | --- | --- | --- | --- |
| 111838673 | *Kv channel-interacting protein 1-like* | LOC111838673 | Cation homeostasis | Large penetrations | Regulatory subunit of Kv4/D (Shal)-type voltage-gated rapidly inactivating A-type potassium channels. Regulates channel density, inactivation kinetics and rate of recovery from inactivation in a calcium-dependent and isoform-specific manner. |
| 111843424 | *potassium voltage-gated channel subfamily C member 2-like* | LOC111843424 | Cation homeostasis | Large penetrations | Voltage-gated potassium channel that mediates transmembrane potassium transport in excitable membranes, primarily in the brain. Contributes to the regulation of the fast action potential repolarization and in sustained high-frequency firing in neurons of the central nervous system |
| 111849794 | *extracellular calcium-sensing receptor-like* | LOC111849794 | Cation homeostasis | Small penetrations | G-protein-coupled receptor that senses changes in the extracellular concentration of calcium ions and plays a key role in maintaining calcium homeostasis. The activity of this receptor is mediated by a G-protein that activates a phosphatidylinositol-calcium second messenger system |
| 111853690 | *protein kinase cGMP-dependent 1* | *prkg1* | Cation homeostasis | Small penetrations | Serine/threonine protein kinase. Numerous protein targets for PRKG1 phosphorylation are implicated in modulating cellular calcium. Proteins that are phosphorylated by PRKG1 regulate platelet activation and adhesion, smooth muscle contraction, cardiac function, gene expression. |
| 111833185 | *myosin XVB* | *myo15b* | Cytoskeletal & sarcomeric | Large penetrations | Unknown, due to the absence of a functional motor domain |
| 111837476 | *troponin C, slow skeletal and cardiac muscles* | LOC111837476 | Cytoskeletal & sarcomeric | Large penetrations | Troponin is the central regulatory protein of striated muscle contraction. The binding of calcium troponin abolishes its inhibitory action on actin filaments |
| 111837614 | *spindle and kinetochore associated complex subunit 3* | *ska3* | Cytoskeletal & sarcomeric | Large penetrations | Component of the SKA1 complex, which is a direct component of the kinetochore-microtubule interface and directly associates with microtubules as oligomeric assemblies |
| 111847443 | *cysteine and glycine rich protein 3* | *csrp3* | Cytoskeletal & sarcomeric | Large penetrations | Positive regulator of myogenesis. Plays a crucial and specific role in the organization of cytosolic structures in cardiomyocytes. It is essential for calcineurin anchorage to the Z line. Can directly bind to actin filaments. Isoform 2 may play a role in early sarcomere organization. |
| 111848724 | *protein tilB homolog* | LOC111848724 | Cytoskeletal & sarcomeric | Large penetrations | May play a role in dynein arm assembly |
| 111850126 | *tubulin beta 2A class Iia* | *tubb2a* | Cytoskeletal & sarcomeric | Large penetrations | Tubulin is the major constituent of microtubules |
| 111852410 | *heat shock protein HSP 90-alpha 1* | LOC111852410 | Cytoskeletal & sarcomeric | Large penetrations | Plays a key role in slow and fast muscle development in the embryo. Plays a role in myosin expression and assembly |
| 111853965 | *tubulin beta chain* | LOC111853965 | Cytoskeletal & sarcomeric | Large penetrations | Tubulin is the major constituent of microtubules |
| 111859691 | *fibroblast growth factor 13* | LOC111859691 | Cytoskeletal & sarcomeric | Large penetrations | Microtubule-binding protein which directly binds tubulin and is involved in both polymerization and stabilization of microtubules |
| 111834243 | *desmin-like* | LOC111834243 | Cytoskeletal & sarcomeric | Small penetrations | Muscle-specific type III intermediate filament essential for proper muscular structure and function. Plays a crucial role in maintaining the structure of sarcomeres. May act as a sarcomeric microtubule-anchoring protein |
| 111856797 | *myosin light chain 3-like* | LOC111856797 | Cytoskeletal & sarcomeric | Small penetrations | Regulatory light chain of myosin. Does not bind calcium. |
| 111833641 | *hyaluronidase-5-like* | LOC111833641 | Extracellular matrix | Small penetrations | Catalyzes the hydrolysis of hyaluronan into smaller oligosaccharide fragments |
| 111848653 | *collagen alpha-1(I) chain-like* | LOC111848653 | Extracellular matrix | Small penetrations | Type I collagen is a member of group I collagen (fibrillar forming collagen) |
| 111857875 | *collagen alpha-1(X) chain-like* | LOC111857875 | Extracellular matrix | Small penetrations | Type X collagen is a product of hypertrophic chondrocytes and has been localized to presumptive mineralization zones of hyaline cartilage |
| 111833176 | *phospholipid-transporting ATPase 1A* | LOC111833176 | Lipid metabolism | Large penetrations | Involved in the transport of aminophospholipids from the outer to the inner leaflet of various membranes and ensures the maintenance of asymmetric distribution of phospholipids |
| 111833450 | *low-density lipoprotein receptor-like* | LOC111833450 | Lipid metabolism | Large penetrations | Binds LDL, the major cholesterol-carrying lipoprotein of plasma, and transports it into cells by endocytosis |
| 111844857 | *ectonucleotide pyrophosphatase/phosphodiesterase 2* | *enpp2* | Lipid metabolism | Large penetrations | Hydrolyzes lysophospholipids to produce the signaling molecule lysophosphatidic acid (LPA) in extracellular fluids. Acts as an angiogenic factor by stimulating migration of smooth muscle cells and microtubule formation. |
| 111845574 | *phospholipase A2-like* | LOC111845574 | Lipid metabolism | Large penetrations | PA2 catalyzes the calcium-dependent hydrolysis of the 2-acyl groups in 3-sn-phosphoglycerides, this releases glycerophospholipids and arachidonic acid that serve as the precursors of signal molecules. |
| 111847497 | *peroxiredoxin-6-like* | LOC111847497 | Lipid metabolism | Large penetrations | It has phospholipase activity |
| 111855292 | *long-chain-fatty-acid--CoA ligase 4-like* | LOC111855292 | Lipid metabolism | Large penetrations | Activation of long-chain fatty acids for both synthesis of cellular lipids, and degradation via beta-oxidation |
| 111853114 | *alkaline ceramidase 2-like* | LOC111853114 | Lipid metabolism | Small penetrations | Hydrolyzes the sphingolipid ceramide into sphingosine and free fatty acid |

DEG = differentially expressed genes, EOD = electric organ discharge.

**Table 6.** Selected DEG in Set C for waveform complexity, by “general” functional class and EOD phenotype, and highlights of their expected function.

| Pking_Entrez_geneID | Pking_gene_name | Pking_gene_symbol | “general” functional class | upregulated in phenotype | Highlights of Predicted Function (edited from UniProt) |
| --- | --- | --- | --- | --- | --- |
| 111838181 | *solute carrier family 9 member A7* | *slc9a7* | Cation homeostasis | Biphasic | Protein: Sodium/hydrogen exchanger 7. Gene: SLC9A7. Mediates electroneutral exchange of protons for Na<sup>+</sup> and K<sup>+</sup> across endomembranes |
| 111838015 | *chloride intracellular channel protein 2-like* | LOC111838015 | Cation homeostasis | Triphasic | Can insert into membranes and form chloride ion channels. Inhibits calcium influx |
| 111848312 | *voltage-dependent calcium channel gamma-1 subunit-like* | LOC111848312 | Cation homeostasis | Triphasic | Regulatory subunit of the voltage-gated calcium channel that gives rise to L-type calcium currents in skeletal muscle. Regulates channel inactivation kinetics |
| 111845832 | *5'-AMP-activated protein kinase subunit gamma-2-like* | LOC111845832 | Cytoskeletal & sarcomeric | Biphasic | AMP/ATP-binding subunit of AMP-activated protein kinase (AMPK). Acts as a regulator of cellular polarity by remodeling the actin cytoskeleton; probably by indirectly activating myosin |
| 111850616 | *transmembrane protein 47-like* | LOC111850616 | Cytoskeletal & sarcomeric | Biphasic | Regulates cell junction organization in epithelial cells. May regulate F-actin polymerization |
| 111851223 | *FYVE, RhoGEF and PH domain-containing protein 4-like* | LOC111851223 | Cytoskeletal & sarcomeric | Biphasic | Plays a role in regulating the actin cytoskeleton and cell shape |
| 111857398 | *protein-methionine sulfoxide oxidase mical2b-like* | LOC111857398 | Cytoskeletal & sarcomeric | Biphasic | Promotes depolymerization of F-actin |
| 111847443 | *cysteine and glycine rich protein 3* | *csrp3* | Cytoskeletal & sarcomeric | Triphasic | Positive regulator of myogenesis. Plays a crucial and specific role in the organization of cytosolic structures in cardiomyocytes. It is essential for calcineurin anchorage to the Z line. Can directly bind to actin filaments. Isoform 2 may play a role in early sarcomere organization |
| 111854588 | *capping protein regulator and myosin 1 linker 3* | *carmil3* | Cytoskeletal & sarcomeric | Triphasic | No info for CARMIL3, but CARMIL2 is a cell membrane-cytoskeleton-associated protein that plays a role in the regulation of actin polymerization at the barbed end of actin filaments. Enhances actin polymerization |
| 111841398 | *inter-alpha-trypsin inhibitor heavy chain 3* | *itih3* | Extracellular matrix | Biphasic | May act as a carrier of hyaluronan in serum or as a binding protein between hyaluronan and other matrix proteins |
| 111841399 | *inter-alpha-trypsin inhibitor heavy chain H3-like* | LOC111841399 | Extracellular matrix | Biphasic | May act as a carrier of hyaluronan in serum or as a binding protein between hyaluronan and other matrix proteins |
| 111853010 | *ependymin-like* | LOC111853010 | Extracellular matrix | Triphasic | GO MF: calcium ion binding. GO BP: cell-matrix adhesion |
| 111853027 | *ependymin-like* | LOC111853027 | Extracellular matrix | Triphasic | GO MF: calcium ion binding. GO BP: cell-matrix adhesion |
| 111853814 | *epiphycan-like* | LOC111853814 | Extracellular matrix | Triphasic | May have a role in bone formation and also in establishing the ordered structure of cartilage through matrix organization |

BP = biological process, DEG = differentially expressed genes, EOD = electric organ discharge, GO = gene ontology, MF = molecular function

### Waveform Duration

Several researchers have implicated the role of ion channels in the evolution of duration changes in mormyrid signals [53–57]. We did not find evidence of large changes in expression of potassium or sodium channels between short-duration *P.* sp. ‘SN3’ and other *Paramormyrops* species. We note that, while no differential expression of potassium or sodium channels was discovered, the GO term ‘regulation of voltage-gated calcium channel activity’ was enriched in long EOD phenotypes, although there were no genes annotated to this GO term common to the lists of DEGs from Sets A’ and B. In particular, individuals with short EODs upregulate two calcium-binding proteins: *parvalbumin-2*, and *parvalbumin-2-like*. Parvalbumins are highly expressed in skeletal muscle where they sequester calcium after contraction, thus facilitating relaxation. Frequently, muscles with fast relaxation rates express higher levels of parvalbumins [76]. The upregulated parvalbumin genes we detected may somehow be related to shorter EODs by sequestering calcium at a faster rate, which could affect action potentials directly or indirectly through calcium-activated ion channels.

Previous studies have demonstrated that changes in EOD duration result from changes in electrocyte ultrastructure. The two major phases of the EOD waveform are caused by action potentials generated by the anterior and posterior faces [47]. Bennett [48] demonstrated a relationship between EOD duration and increased surface membrane area, and Bass et al. [50] showed that differences in surface area are more readily noticeable on the anterior face. Membrane surface area is increased by folding the electrocyte membrane into papillae and other tube-like invaginations [77]. Testosterone can induce increases in EOD duration in several mormyrids [49,50,78,79], and it also increases membrane surface area, either particularly on the anterior face [50] or on both anterior and posterior faces [80]. A larger surface area may increase the capacitance of the membrane, thus delaying spike initiation [49,50]. Consequently, genes involved in the synthesis of membranes could influence EOD duration.

We found the most prominent differences in gene expression between the EOD duration phenotypes in genes that code for cytoskeletal, sarcomeric, and lipid metabolism proteins (Table 4). We emphasize the last group: no lipid metabolism genes were upregulated in individuals with long EODs, whereas samples with short EODs upregulated *protein EFR3 homolog B-like* (a regulator of phosphatidylinositol 4-phosphate synthesis), *retinoic acid receptor responder protein 3-like* (hydrolysis of phosphatidylcholines and phosphatidylethanolamines), *PTB domain-containing engulfment adapter protein 1-like* (modulates cellular glycosphingolipid and cholesterol transport), *phosphatidylinositol 3-kinase regulatory subunit gamma-like*, (PI3K, which phosphorylates phosphatidylinositol), and *proto-oncogene c-Fos-like* (can activate phospholipid synthesis), and showed enrichment of the GO term ‘phosphatidylinositol phosphorylation.’ We hypothesize that these genes are involved in the surface proliferation of the electrocytes membranes.

Additionally, each mormyrid electrocyte stands embedded in a gelatinous mucopolysaccharide matrix (the ECM) separated from neighboring electrocytes by connective tissue septa (Fig. 1) [34], and the membrane surface invaginations are coated by the same ECM that surrounds the electrocytes [50,77]. Hence, differences in surface invaginations could also be reflected in differences in the expression of genes whose products interact with the ECM. We detected none of these genes upregulated in individuals with long EODs, whereas those with short EODs upregulated three: *hyaluronidase-5-like* (breaks down hyaluronan), *collagenase 3-like* (plays a role in the degradation of ECM proteins), and *fibroblast growth factor binding protein 1* (acts as a carrier protein that releases fibroblast-binding factors from the ECM storage).

Overall, our results identify genes that may affect EOD duration through membrane rearrangements, which could be coupled with changes in the interaction with the ECM and the expression of cytoskeletal and sarcomeric genes. Since this waveform feature is modulated by testosterone, this androgen could facilitate the study of these suggested genetic underpinnings under more rigorously controlled circumstances.

### Waveform Polarity

The number [47] and diameter [34,43] of stalk penetrations are positively correlated with the magnitude of P0. This phenomenon is exemplified by *P.* sp. ‘magnostipes type I’, which has the largest P0 in the OTUs examined in this study, giving the EOD the appearance that it ‘inverted’ relative to other EODs. This OTU has numerous, large diameter penetrations, whereas *P. kingsleyae* (P-type) has relatively fewer, small diameter penetrations (Fig. 1). These large structural differences may influence the electrocyte’s connection with the surrounding ECM, and our results support this: the phenotypes of waveform polarity exhibited differences in the expression of genes that interact with the extracellular space. We found no such genes upregulated in individuals with large penetrations, whereas in samples with small penetrations two GO terms were enriched: ‘extracellular matrix organization’ and ‘response to mechanical stimulus,’ and three genes were upregulated: *collagen alpha-1(I) chain-like*, *collagen alpha-1(X) chain-like*, and *hyaluronidase-5-like* (breaks down hyaluronan).

OTUs with large penetrations also exhibited higher expression of genes related to cytoskeletal, sarcomeric, and lipid metabolism proteins than do individuals with smaller penetrations (Table 5). This includes the GO terms ‘heart contraction,’ ‘sarcomerogenesis,’ and ‘cardiac myofibril assembly,’ representative of the genes *myosin XVB*, *heat shock protein HSP 90-alpha 1* (has a role in myosin expression and assembly), *tubulin beta chain*, *tubulin beta 2A class IIa*, *fibroblast growth factor 13* (a microtubule-binding protein which directly binds tubulin and is involved in both polymerization and stabilization of microtubules), *spindle and kinetochore associated complex subunit 3* (a component of the kinetochore-microtubule interface), *troponin C, slow skeletal and cardiac muscles*, *protein tilB homolog* (may play a role in dynein arm assembly), and *cysteine and glycine rich protein 3* (codes for the Muscle LIM Protein (MLP), which is implicated in various cytoskeletal and sarcomeric macromolecular complexes [81–83] and is a positive regulator of myogenesis [84]). In contrast, samples with small penetrations only show upregulation of the genes *myosin light chain 3-like* and *desmin-like*.

We hypothesize that the differences in the number and diameter of penetrations that drive variation in EOD waveform polarity require changes to the electrocyte’s cytoskeletal and membrane properties. These arrangements may be necessary for the electrocytes body to adjust to the increased volume displacements imposed by larger penetrations; or alternatively, they may be a prerequisite for penetrating stalks to enlarge. Our observations support and elaborate on the hypothesis that sarcomeric proteins (which are non-contractile in mormyrids) may function as a means of cytoskeletal support and structural integrity in mormyrid electrocytes [85].

### Waveform Complexity

Waveform complexity refers to the number of phases present in an EOD, and mormyrid EODs vary in the presence of a small head negative phase (P0). The presence or absence of P0 in the EOD depends on the anatomical configuration of the electrocytes: P0-present (or triphasic) EODs are produced by electrocytes that are innervated on the anterior face and have penetrating stalks (Pa), whereas P0-absent (or biphasic) EODs are produced by electrocytes innervated on the posterior face and lack penetrating stalks (NPp) [42,43,47,48,51,52]. Developmental studies of the adult EO suggest that Pa electrocytes go through a NPp stage before developing penetrations [86,87]. This motivated the hypothesis that penetrations develop by the migration of the posteriorly innervated stalk system (NPp stage) through the edge of the electrocyte, and that the interruption of this migration represents a mechanism for Pa-to-NPp reversals [88,89].

Our data indicates several DEGs that implicate specific cytoskeletal and ECM reorganizations between triphasic and biphasic EODs (Table 6). We observed differential expression of several genes associated with the polymerization of F-actin. In triphasic individuals, we observe upregulation of the gene *capping protein regulator and myosin 1 linker 3* (CARMIL3); although this gene is little studied, its paralog CARMIL2 enhances F-actin polymerization. Also upregulated is the gene *cysteine and glycine rich protein 3* (MLP protein, see Waveform Polarity). In contrast, the biphasic phenotype upregulated the genes *protein-methionine sulfoxide oxidase mical2b-like* (promotes F-actin depolymerization), *transmembrane protein 47-like* (may regulate F-actin polymerization), 5’-*AMP-activated protein kinase subunit gamma-2-like* (could remodel the actin cytoskeleton), and *FYVE, RhoGEF and PH domain-containing protein 4-like* (regulates the actin cytoskeleton). Thus, biphasic and triphasic EODs display several DEG, with potentially diverging outcomes, that influence the cellular internal structure.

We hypothesize that electrocytes with penetrating stalks (which produce triphasic EODs) require cytoskeletal arrangements to produce penetrations, perhaps related to increasing F-actin, to maintain their structural integrity. Similar to what we propose under waveform polarity, these arrangements may be necessary for the electrocyte body to adjust to the penetrations; or alternatively, they may be a prerequisite for penetrations to occur.

We also observed differential expression in a number of proteins expressed in the ECM. In biphasic OTUs, we found the GO term ‘mucopolysaccharide metabolic process’ to be enriched, and two upregulated copies of the gene *inter-alpha-trypsin inhibitor heavy chain 3*, which may act as a binding protein between hyaluronan and other ECM proteins. In triphasic individuals, we found the GO term ‘cell-matrix adhesion’ enriched, and the upregulated genes *epiphycan-like*, which may play a role in cartilage matrix organization, and two copies of *ependymin-like* (these paralogs are ortholog to the zebrafish ependymin-like gene *epdl2*).

The two ependymin-like genes are among the most differentially expressed genes between biphasic and triphasic OTUs (500-fold more highly expressed in triphasic individuals in the comparison from Set A’, Supplemental File 2). Although expressed in many tissues and with little amino acid similarity, all ependymin-related proteins are secretory, calcium-binding glycoproteins that can undergo conformational changes and associate with collagen in the ECM. They have been involved in regeneration, nerve growth, cell contact, adhesion and migration processes [90]. We hypothesize that ependymin-related proteins, and potentially some of the other ECM proteins highly expressed in triphasic individuals, are part of the “fibrillar substance” that lies between the stalk and the electrocyte body in individuals with penetrating electrocytes [50]. Notably, the *P. kingsleyae* genome assembly, which is based on a biphasic individual, contains three paralogs of *epdl2*, whereas the osteoglossiform *Scleropages formosus* only has one, suggesting the intriguing possibility that this gene may have been duplicated in *Paramormyrops* or in mormyrids. Ependymin-related paralogs have been proposed as suitable targets to experimentally test gene subfunctionalization [91].

Altogether, our results for EOD waveform complexity suggest that the conformation of the cytoskeleton and the expression of proteins secreted to the ECM are important elements of the stalk penetrations, which generate triphasic EODs.

### Concluding Remarks

Two previous studies focused on DGE between EOs in another mormyrid species adaptive radiation/explosive diversification (genus *Campylomormyrus*). Both focused on comparisons between the species *C. tshokwe* (long duration) and *C. compressirostris* (short duration). The first study performed a canditate gene approach to quantify the expression patterns of 18 sodium and potassium homeostasis genes between the EOs of the two species [55], whereas Lamanna et al. [92] used RNA-seq to simultaneously compare gene expression between EOs of these species. While we did not observe differences in expression of any of the potassium channels reported by Nagel et al. (2017), we note that Lamanna et al. (2015) reported differential expression of metabolic pathways related genes, particularly fatty acid metabolism, and ion transport and neuronal function (subcluster 4). While we found no overlap in the identities of any specific genes in our study, we note that our analysis also detected differential expression of lipid metabolism related genes when comparing EODs of different duration.

The widespread differential expression within *Paramormyrops* of calcium-related genes (Supplemental File 6) emphasizes a much-needed area of future research. Calcium is known to be necessary for the proper electrocyte repolarization in some gymnotiform species [93], but it may not be as important in others [94]. Few studies have addressed calcium physiology in mormyrids: calcium-related proteins have been reported as differentially expressed in EO vs skeletal muscle in *Campylomormyrus* [92] and in *Brienomyrus brachyistius* [85]. As electrocytes do not contract, calcium may act in electrocytes as an important second messenger or cofactor, participate in interactions with the ECM, and/or to contribute to the electrocyte’s electrical properties through interaction with voltage gated ion channels.

A second notable pattern in our results is the unusual degree to which mormyrid electrocytes retain expression of some sarcomeric genes, which has been noted in several studies [58,85,92,95,96]. The role these proteins serves in electrocytes is presently unknown; however, results indicate that they are highly differentially expressed between *Parmormyrops* with different EOD waveforms. This strongly suggests that sarcomeric proteins could play an important role in the conformational changes required to develop and sustain penetrations.

Finally, the biochemical composition and function of the ECM in electrocytes is poorly understood. Our analysis identifies differential expression in ECM-related genes across the *Paramormyrops*, associated with each of the three EOD features studied. At least two of these genes (*inter-alpha-trypsin inhibitor heavy chain 3* and *hyaluronidase-5-like*), distributed across all three EOD features, interact with hyaluronan. Hyaluronan is a type of mucopolysaccharide and a major component of some soft tissues and fluids [97]. Therefore, we propose that hyaluronan is an important constituent of the ECM in mormyrid fish. In addition, the electrocyte-ECM interactions should be an important area of future investigation, as they are likely to influence electrocyte shape, electrical properties, and potentially the morphology of penetrations and surface membrane invaginations.

To conclude, this study examined the expression correlates of a hyper-variable phenotype in a rapidly diversified genus of mormyrid electric fish. We examined DGE between taxa exhibiting variability along three major axes of variation that characterize EOD differences within *Paramormyrops* and among mormyrids: duration, polarity, and complexity. We found that gene expression in EOs among closely related species is largely similar, but patterns of DGE between EOs is primarily restricted to four broad functional sets: (1) cytoskeletal and sarcomeric proteins, (2) cation homeostasis, (3) lipid metabolism and (4) proteins that interact with the ECM. Our results suggest specific candidate genes that are likely to influence the size, shape and architecture of electrocytes for future research on gene function and molecular pathways that underlie EOD variation in mormyrid electric fish.

## Ethics approval and consent to participate

All research protocols involving live fish were approved by the Michigan State University and Cornell University Institutional Animal Care and Use Committees.

## Acknowledgements

This work was supported by the National Science Foundation (1455405: JRG, PI) and a grant from the Cornell University Center for Vertebrate Genomics. The authors thank Bruce Carlson, Matt Arnegard, Carl Hopkins, Roger Afene, Jean Danielle Mbega and Marie-Francois Eva for assistance with specimen collection in Gabon. The authors also acknowledge the Michigan State University Institute for Cyber-Enable Research for use of their high-performance computing infastructure, and CENREST Gabon for logistical support and collection permits.

## Author Contributions

JRG designed the experiment, collected and identified specimens, performed library construction and sequencing, and oversaw data analysis. ML performed QC analysis, designed and implemented the data analysis procedure, and performed examination of gene ontology and gene function. Both authors contributed to writing the manuscript.

## Supplementary Files

**Supplementary File 1.** Raw reads NCBI SRA accession numbers, number of reads and alignment rates per sample, using bowtie 2 as the aligner and the Paramormyrops kingsleyae (N-type) genome as the reference.

**Supplementary File 2.** DEG per comparison from the 10 pairwise DGE analysis. Positive values under logFC indicate genes upregulated in the OTU under sampleA, whereas negative values correspond to genes upregulated in the OTU under sampleB. Values under each sample are gene raw counts. Significance threshold was abs(log(base2)FC) > 2 (= 4-fold expression difference) and FDR <0.001.

**Supplementary File 3.** Enriched GO terms per comparison, ontology and OTU in the DEG from the 10 pairwise comparisons. Also listed are the DEG annotated to each GO term. The pvalue is in the column weight01.

**Supplementary File 4.** DEG per EOD feature and phenotype identified with the Set B analysis. Values under each sample are TMM normalized, log2(TMM +1) transformed, and mean-centered expression values.

**Supplementary File 5.** Enriched GO terms per EOD feature, ontology and phenotype in the DEG from the Set B analysis. Also listed are the DEG annotated to each GO term. The pvalue is in the column weight01.

**Supplementary File 6.** DEG in Set C, per EOD feature and phenotype.

**Supplementary File 7.** GO terms enriched in the DEG in Set C, per EOD feature, ontology and phenotype. Also listed are the DEG annotated to each GO term, and the quickGO definitions of each GO term.

